# Mount Fuji’s stubby peak: the genotypic density of additive landscapes near maximal fitness

**DOI:** 10.64898/2026.04.02.716185

**Authors:** Justin B. Kinney

## Abstract

Additive fitness landscapes—also called Mount Fuji landscapes—are the simplest and most widely used models of sequence-function relationships. As such, they play essential roles across multiple areas of biology, including evolutionary theory, quantitative genetics, gene regulation, and protein science. One of the most basic properties of any fitness landscape is its *genotypic density*—the number of sequences near a given fitness value. Understanding this density is especially important near fitness peaks, as it quantifies the supply of high-fitness genotypes. Here I study the genotypic density of additive landscapes near fitness peaks. Although this density is well known to be approximately Gaussian near the middle of the fitness range, its behavior near maximal fitness has not been reported. I begin by deriving a saddle-point approximation that accurately describes the genotypic density of additive landscapes over virtually the entire fitness range. I then show that the log density follows a power law near maximal fitness, with the exponent determined by how much the best allele at each position outperforms its nearest competitor. This power-law behavior holds over a substantial fraction of fitness values, besting the Gaussian approximation on both simulated and empirical landscapes across roughly a quarter to a third of the fitness range. Under certain conditions this behavior also extends to globally epistatic landscapes (defined as nonlinear functions of one or more additive traits), though with a reduced range of validity. These findings advance our understanding of one of the most fundamental models of sequence-function relationships. In particular, they reveal that the uppermost reaches of Mount Fuji landscapes, rather than being sharply peaked, are actually quite stubby.

## Introduction

Fitness landscapes—quantitative mappings from biological sequences to fitness values^1^—have long been a key organizing concept in evolutionary biology (Wright 1932; Maynard Smith 1970; Fragata et al. 2019). The simplest and most widely used model for such landscapes is the *additive model*, in which each position in a sequence contributes independently to fitness. Additive models are used in many areas of biology, both in theoretical efforts and in empirical studies of sequence-function relationships. In particular, they form the basis for *global epistasis models*, which are defined as nonlinear functions of one or more additive traits. Many experimentally measured fitness landscapes are well-described by either additive or globally epistatic models (Carneiro and Hartl 2010; Kinney et al. 2010; Kryazhimskiy et al. 2014; Stiffler et al. 2015; Sarkisyan et al. 2016; Adams *et al*. 2016; Sailer and Harms 2017; Otwinowski 2018; Otwinowski *et al*. 2018; Baeza-Centurion et al. 2019; Starr et al. 2020; Faure *et al*. 2024). A thorough understanding of the quantitative properties of additive models is therefore warranted.

One of the simplest questions one can ask about a fitness landscape is how many sequences have fitness near a given value. I refer to this quantity as *genotypic density*. In the case of additive landscapes, the Central Limit Theorem (CLT) suggests that the genotypic density is approximately Gaussian in the bulk of the fitness distribution. However, this bulk approximation is qualitatively wrong near the maximal achievable fitness. Although the importance of genotypic density has been recognized by many authors, both for modeling evolution (Berg and von Hippel 1987; Aita and Husimi 1996; Sengupta *et al*. 2002; Djordjevic *et al*. 2003; Berg *et al*. 2004; Mustonen and Lassig 2005; Sella and Hirsh 2005; Lässig 2007; Mustonen *et al*. 2008; Neher and Shraiman 2011) and for bioinformatic analyses (Staden 1989; Touzet and Varré 2007; Madsen *et al*. 2017), the quantitative form of the near-peak density of additive fitness landscapes has not been reported.

In the fitness landscape literature, additive landscapes are sometimes known as *Mount Fuji* landscapes, a moniker chosen to emphasize their simple topographic structure: a single peak that is evolutionarily accessible from any starting sequence (Aita and Husimi 1996, 2000; Aita *et al*. 2004). Although the Gaussian behavior of genotypic density in the bulk was realized early on in this context (Aita and Husimi 1996), the behavior near maximal fitness appears not to have been investigated. This is perhaps because additive landscapes have primarily been considered as null models against which to assess more complex epistatic models (including NK, Rough Mount Fuji, and House of Cards models) (Szendro *et al*. 2013; Neidhart *et al*. 2014; Pahujani and Krug 2025), rather than as objects of primary scientific interest.

The near-peak genotypic density nevertheless has evolutionary implications: it quantifies the supply of high-fitness genotypes accessible by mutation, and therefore shapes both adaptive and purifying selection.

In quantitative genetics, additive models have long provided a theoretical foundation for the study of plant and animal breeding (Walsh and Lynch 2018; Barton and Keightley 2002). Of particular importance is the infinitesimal model, which assumes that a trait is determined by the sum of effects across a large number of loci (Barton et al. 2017). Both theory and data indicate that additive genetic contributions account for most of the heritable variation in complex traits, even when epistasis is present at individual loci (Hill et al. 2008). The distribution of trait values in a population under selection has been studied extensively, including using concepts and methods from statistical physics (Barton and Coe 2009; Barton and de Vladar 2009). In particular, the departure of population trait distributions from Gaussianity has been explored (Turelli and Barton 1994). Genotypic density itself, however, does not appear to have received a similar treatment.

In the bioinformatic study of gene regulation, additive landscapes are commonly used to model the binding of transcription factors (TFs) to DNA. Additive models such as position weight matrices (PWMs) are routinely used to scan genomes for putative TF binding sites (Stormo 2000; Stormo and Fields 1998; Foat et al. 2006; Djordjevic et al. 2003; Kinney et al. 2007; Elemento et al. 2007; Grant *et al*. 2011), and large databases of such models are widely used (Baydar et al. 2025; Vorontsov et al. 2024; Wingender et al. 2000; Pachkov et al. 2007). Assessing the statistical significance of high-scoring sites requires accurate estimates of the upper tail of the genotypic density, and a few studies have developed methods for doing so (Staden 1989; Touzet and Varré 2007; Madsen et al. 2017). Of particular note is Madsen *et al*. (2017), who developed an estimate of p-values for PWMs using a saddle-point approximation similar to the one I use below. However, neither Madsen *et al*. (2017) nor other work in this area has described the mathematical form of the near-peak density.

Beginning with foundational studies by Berg and von Hippel (Berg and von Hippel 1987, 1988), researchers at the interface of statistical physics and population genetics have used additive models for TF binding energy to study the evolution of TF binding sites (Sella and Hirsh 2005; Sengupta et al. 2002; Gerland *et al*. 2002; Djordjevic et al. 2003; Neher and Shraiman 2011). Genotypic density plays an important role in this literature. When evolution is modeled in the weak-mutation limit, the equilibrium distribution of binding site sequences takes the form of a canonical ensemble from statistical physics, one in which fitness values are weighted by an exponentially tilted form of the genotypic density (Berg et al. 2004; Mustonen and Lassig 2005; Mustonen et al. 2008; Lässig 2007). The original work of Berg and von Hippel derives the exact genotypic density for an idealized additive model in which every mutation away from the highest-fitness sequence produces the same fitness deficit. To treat more general additive models, Sengupta *et al*. (2002) derived a saddlepoint estimate of genotypic density (the same estimate I consider below), but used it only to recover the Gaussian approximation suggested by the CLT.

Here I study the genotypic density of additive landscapes near maximal fitness. I first derive a saddle-point approximation to this density and show that it is remarkably accurate across nearly the entire fitness range. I then investigate the near-peak behavior of this approximation. I find that, for an important class of idealized models, the log density follows a power law (or equivalently, the density itself has an exponentiated powerlaw form). Additional considerations as well as empirical results suggest that many real-world landscapes also follow an exponentiated power law. A consequence is that, as fitness decreases from its maximum, the number of available genotypes initially expands very rapidly, then grows more and more slowly until finally it matches up with the predictions of the Gaussian approximation. The landscape is therefore not sharply peaked at its maximum but instead is broad and gently rounded—more like the top of a rolling hill than the summit of a mountain. This motivates my use of the term “stubby.”

I validate these findings on simulated landscapes and on two empirical landscapes: one from a massively parallel reporter assay on a transcriptional promoter (Kinney *et al*. 2010), and one from a deep mutational scanning experiment on a protein domain (Olson *et al*. 2014). In all cases these landscapes exhibit stubby peaks that are well described by an exponentiated power law. I then show that globally epistatic landscapes defined on one or more traits often exhibit near-peak densities of the same mathematical form, though with caveats. The derivations of these results are provided below. Additional mathematical details are given in Supplemental Information (SI).

## Results

### Preliminaries

Consider sequences of length *L* comprising characters from an alphabet of size *C* (e.g. *C* = 4 for DNA or RNA, *C* = 20 for proteins). Each sequence *x* is represented by a one-hot encoded vector *x* = {*x*_*lc*_}, where *x*_*lc*_ = 1 if character *c* occurs at position *l* and *x*_*lc*_ = 0 otherwise. Additive fitness landscapes have the form

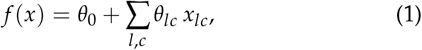

where *θ*_*lc*_ quantifies the effect of character *c* at position *l* and *θ*_0_ is an overall fitness baseline. In what follows I denote the maximal fitness effect at position *l* by 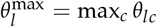, the optimal character by 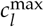, and the fitness-maximizing sequence by *x*_max_. The maximum achievable fitness is *F*_max_ = *f* (*x*_max_). *F*_min_ denotes minimum achievable fitness.

Throughout this paper I illustrate the claims and findings using additive models for two empirical landscapes (Fig. 1). The first model describes the transcriptional activity of variants of the *Escherichia coli lac* promoter (*L* = 75, *C* = 4) measured using a massively parallel reporter assay (MPRA) called Sort-Seq (Kinney *et al*. 2010). The second model describes the binding of variants of the B1 domain of protein G (GB1; *L* = 55, *C* = 20) to IgG, as measured using a deep mutational scanning (DMS) experiment based on RNA display (Olson *et al*. 2014). Additive models for both landscapes were inferred from experimental data using MAVE-NN (Tareen *et al*. 2022), which separately accounts for any global epistasis nonlinearities present in the data.

**Figure 1.**
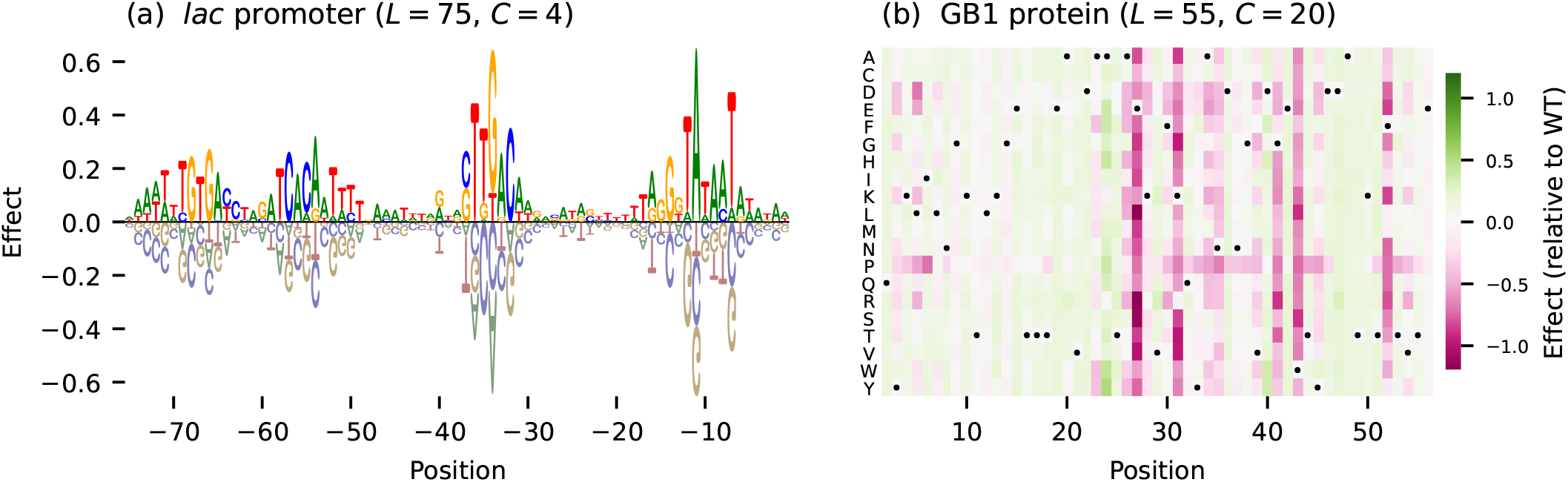
Two empirical landscapes analyzed in this paper. (a) Additive model for the *lac* promoter landscape of Kinney *et al*. (2010) (*L* = 75, *C* = 4), visualized as a sequence logo (Tareen and Kinney 2019). Positions are relative to the transcription start site. (b) Additive model for the GB1 landscape of Olson *et al*. (2014) (*L* = 55, *C* = 20), visualized as a heatmap. Positions correspond to residues 2-56. Dots mark wild-type residues. The parameters of both models were inferred from data using MAVE-NN (Tareen et al. 2022), then shifted and scaled so that the fitness values represent z-scores, i.e., random sequences have fitness values with zero mean and unit variance.

### Defining the genotypic density

The focus of this paper is the quantitative form of the genotypic density, *ρ*(*F*), which is defined so that *ρ*(*F*) *dF* is the number of sequences within an infinitesimal interval [*F, F* + *dF*]. Formally, this density is written in terms of Dirac delta functions:

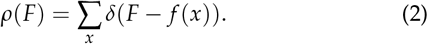

Below I develop various smooth mathematical approximations to *ρ*(*F*). The reader might reasonably question what it means to approximate *ρ*(*F*) with a smooth function, since no nonzero smooth function will provide a good approximation in the standard squared-error sense.

Perhaps the most intuitive reason for considering smoothed approximations to *ρ*(*F*) is that, in reality, fitness values are never measured with infinite precision. One way to account for this is to replace the delta function in Eq. 2 with a Gaussian distribution that has a standard deviation commensurate with the expected level of experimentally unresolvable uncertainty. Doing this yields a kernel density estimate that is smooth by construction. In practice, the form of this estimate is almost completely independent of the width of the Gaussian as long as the uncertainty scale is much less than the fitness range *F*_max_ − *F*_min_, and much greater than the gaps between the extremal fitness values (*F*_max_ and *F*_min_) and their nearest neighbors. This condition is satisfied for a wide range of uncertainties, and almost always holds in practice.

### The bulk approximation

In an additive landscape, the fitness of each sequence is a sum of *L* independent contributions. The CLT therefore suggests that *F* values are approximately normally distributed, and thus that *ρ*(*F*) can be approximated by

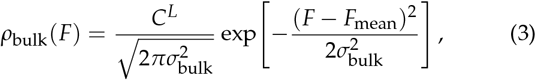

where *F*_mean_ and 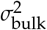are the mean and variance of the fitness values observed for randomly generated sequences. The CLT, however, only guarantees accuracy within a few standard deviations of the mean. Indeed, Fig. 2 shows that this Gaussian form accurately describes the central mass of *ρ*(*F*), but breaks down dramatically near *F*_max_. Specifically, *ρ*_bulk_(*F*) provides increasingly inflated estimates of *ρ*(*F*) as *F* approaches *F*_max_ from below, and predicts nonzero density at all fitness values greater than *F*_max_, where the true density vanishes. The breakdown worsens as sequences grow longer. Assuming the distribution of *θ*_*lc*_ values remains fixed, the width of the fitness distribution scales as 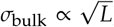, whereas *F*_max_−*F*_mean_ grows linearly with *L*. The fitness peak therefore recedes to ever more extreme quantiles as *L* increases, placing it in a regime where the CLT provides no useful information. To characterize *ρ*(*F*) across the full fitness range—and in particular near *F*_max_—a fundamentally different approach is required.

**Figure 2.**
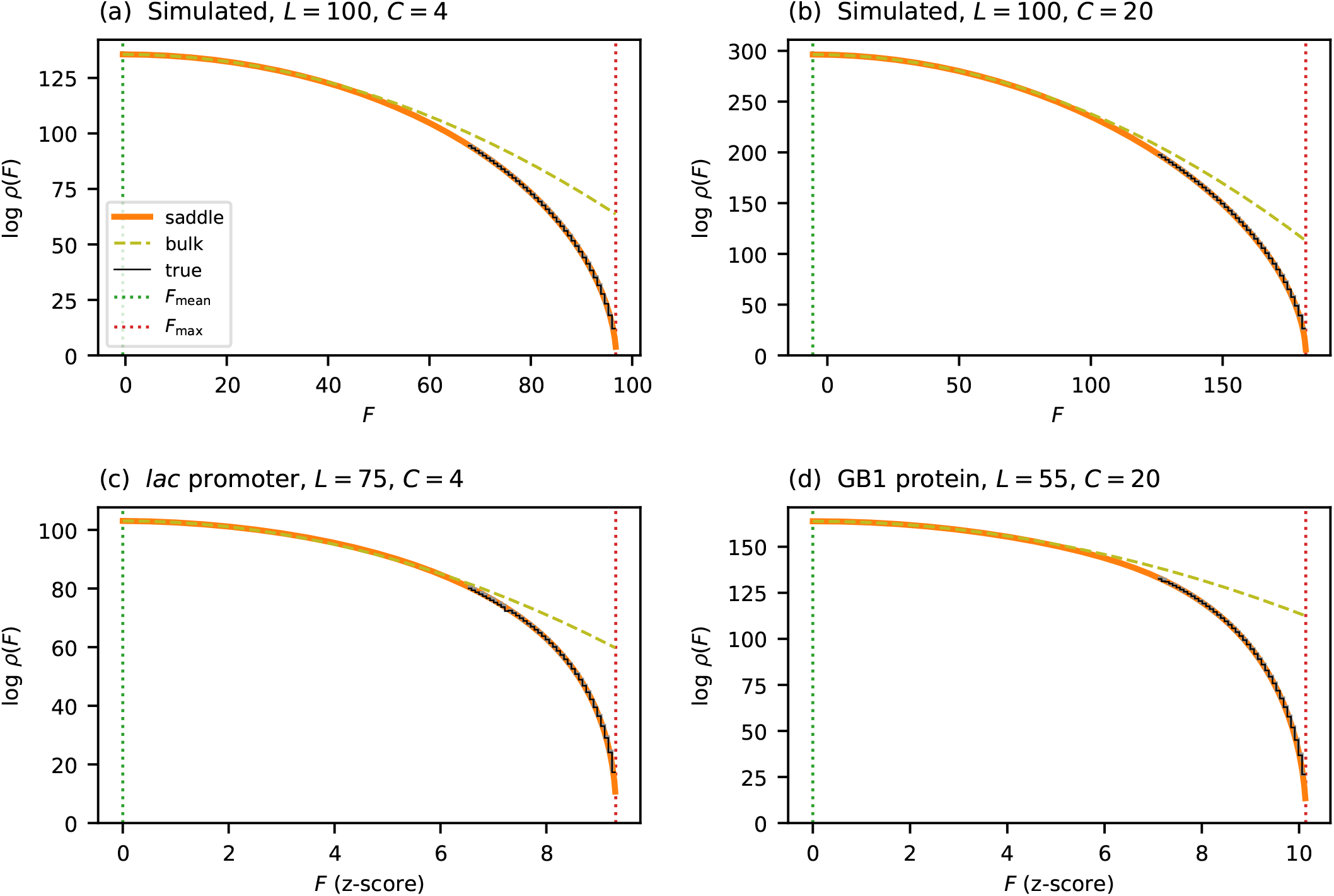
Accuracy of the saddle-point approximation on simulated and empirical landscapes. (a,b) Results for simulated landscapes having Gaussian effects *θ*_*lc*_ and alphabets of different size. (c) Results for the *lac* promoter landscape of Kinney *et al*. (2010). (d) Results for the GB1 landscape of Olson *et al*. (2014). Fitness values for the empirical landscapes correspond to z-scores, as in Fig. 1. Each panel compares the saddle-point approximation *ρ*_saddle_ (orange solid line) to both the Gaussian approximation *ρ*_bulk_ (olive dashed line) and the true density *ρ*_true_ (black solid line), computed using the algorithm of Touzet and Varré (2007). Vertical dotted lines indicate *F*_max_ and *F*_mean_.

### The tilted distribution and free fitness

To derive an improved estimate of *ρ*(*F*), we first introduce a family of *exponentially tilted* distributions over sequence space:

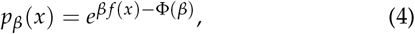

where *β* is a scalar that parameterizes the family, and

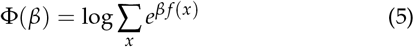

is a function that serves to normalize the distribution. By varying *β* from − ∞ to ∞ we can center this distribution over specific fitness values. In particular, *β* = 0 recovers the uniform distribution on sequence space (which has fitness values concentrated near *F*_mean_), while sufficiently large *β* concentrates the distribution near *F*_max_.

Distributions of the form in Eq. 4 arise naturally in both population genetics and statistical physics. In the weak-mutation limit of Wright-Fisher processes with reversible mutation rates, *p*_*β*_ (*x*) describes the stationary distribution over genotypes, where *β* is related to the effective population size via *β* = 2*N*_*e*_ (Berg *et al*. 2004; Mustonen and Lassig 2005; Lässig 2007; Sella and Hirsh 2005; Neher and Shraiman 2011). If instead we interpret *x* as the microstate of a physical system and − *f* (*x*) as the corresponding energy, then *p*_*β*_ (*x*) describes the distribution over microstates that arises when the system is at temperature 1/*β*. In such contexts *p*_*β*_ is called the *canonical ensemble*.

The normalizing term Φ(*β*) encodes important information about the tilted distribution. Indeed, this quantity has previously been termed *free fitness* (Iwasa 1988; Sella and Hirsh 2005; 4 Mount Fuji’s stubby peak Lässig 2007; Barton and de Vladar 2009), as it is analogous to negative the free energy in physical systems. One key property is that Φ(*β*) is a cumulant-generating function for the marginal distribution of fitness values,

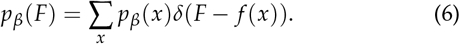

In other words, the mean of *p*_*β*_ (*F*) is *µ*_*β*_ = Φ^′^(*β*), the variance is 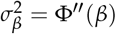, and so on. This is true regardless of the functional form of *f* (*x*).

In the specific case where *f* (*x*) is additive (Eq. 1), free fitness can be written in closed form:

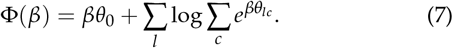

Corresponding expressions for *µ*_*β*_ and 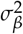 are given in SI Sec. S1. It is readily shown that the higher-order cumulants become irrelevant as *L* grows, and thus that *p*_*β*_ (*F*) becomes well-approximated by the Gaussian distribution

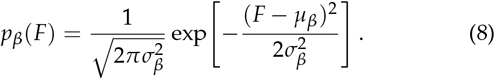

### The saddle-point approximation

We now return to the problem of estimating the genotypic density *ρ*(*F*) in Eq. 2. Since the weight for each sequence in the tilted distribution depends on *x* only through *f* (*x*), it can be taken outside the sum in Eq. 6. Additionally using Eq. 2 and Eq. 4 then gives

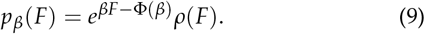

We now have two distinct expressions for *p*_*β*_ (*F*), Eq. 8 and Eq. 9. Equating these and solving for *ρ*(*F*) yields a candidate approximation for genotypic density

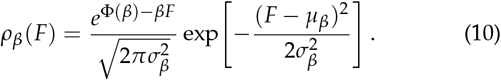

We are still free to choose *β* however we like. We expect, however, that *ρ*_*β*_ (*F*) will be a good approximation only when *F* is within a few standard deviations *σ*_*β*_ of the mean *µ*_*β*_. Indeed, Eq. 10 reduces to the bulk approximation Eq. 3 if we set *β* = 0, and the entire motivation for this paper is that *ρ*_bulk_(*F*) is a poor estimate of the true density *ρ*(*F*) when *F* is near *F*_max_.

The CLT suggests that the approximation in Eq. 8 is most accurate when *µ*_*β*_ = *F*. We therefore choose *β* to be a function of *F* defined implicitly by this requirement, i.e.,

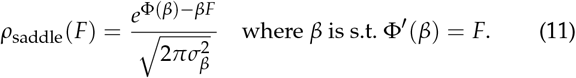

This is known as the *saddle-point* approximation (Darwin and Fowler 1922; Daniels 1954; Reid 1988; Barndorff-Nielsen and Cox 1989). The name comes from an alternative derivation in which one approximates the value of a contour integral in the complex plane by its value near a stationary point of the integrand (i.e., a saddle-point). The above derivation, however, more clearly explains why the saddle-point approximation works so well throughout the entire fitness range. See Reid (1988) for further discussion of these two derivations. Eq. 11 can also be thought of in terms of large deviation theory (Touchette 2009), with rate function given by [*βF* − Φ(*β*)]/*L*. The above derivation, however, has the advantage of also providing the 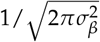prefactor, which is generally neglected in rate function calculations.

### Performance of the saddle-point approximation

To assess the performance of *ρ*_saddle_, I compared its predictions to the true densities of two simulated landscapes and of the two empirical landscapes mentioned earlier. The results are shown in Fig. 2. In all cases *ρ*_saddle_ closely tracks *ρ* over the entire fitness range. By contrast, *ρ*_bulk_ progressively overestimates *ρ* as *F* approaches *F*_max_, ultimately deviating by many orders of magnitude. This agreement holds equally for simulated and empirical landscapes over both DNA (*C* = 4) and protein (*C* = 20) alphabets.

I note that computing the true density *ρ* is complicated by the fact that all four landscapes in Fig. 2 are too large to exhaustively enumerate the fitness values of all sequences. I therefore used the dynamic programming algorithm of Touzet and Varré (2007) instead. This algorithm computes cumulative counts above a given threshold by iteratively refining upper and lower bounds using discretized fitness values. Running this algorithm to full precision across the entire fitness range is prohibitive. Restricting it to the upper 30% of the range and terminating refinement before full convergence, however, yields certified upper and lower bounds that are visually indistinguishable in Fig. 2.

### Near-peak scaling for binary alphabets

Eq. 11 does not by itself reveal the mathematical form of genotypic density near maximal fitness. What we require is an explicit expression for *ρ*(*F*) near *F*_max_, one that is not obscured by the implicit dependence of *β* on *F*. To this end, we express the free fitness as Φ(*β*) = *βF*_max_ + *LE*(*β*), where

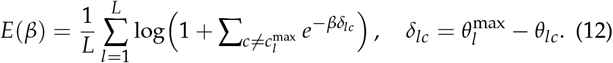

We call this the *per-site deficit contribution* to free fitness, and define the *per-site fitness deficit* as *ϵ* = (*F*_max −_*F*)/*L*. Substituting these into Eq. 11 and dropping the prefactor (which is sub-leading for large *L*) gives

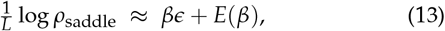

where *β* is an implicit function of *ϵ* determined by *E*′(*β*) = − *ϵ*. Since every *δ*_*lc*_ > 0 in Eq. 12, *E*(*β*) → 0 as *β* → ∞. However, the near-peak behavior of *ρ*_saddle_ depends on precisely how *E*(*β*) approaches this limit. We therefore seek to determine the leading-order behavior of *E*(*β*) as *β* → ∞.

To simplify this analysis, we first consider binary alphabets (*C* = 2). In this case, the deficit contribution becomes

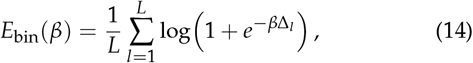

where Δ_*l*_ is the (positive) fitness gap between the two additive effects at position *l*. Replacing the empirical distribution of these gaps with a smooth density *p*_gap_(Δ) gives

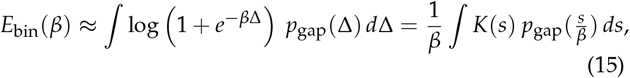

where we have substituted *s* = *β*Δ and defined *K*(*s*) = log(1 + *e*^−*s*^). It is helpful to think of *K*(*s*) as a smoothing kernel: it equals log 2 at the origin, has width of order 1, decays as *e*^−*s*^ for *s* ≫ 1, and has area 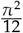 (see SI Sec. S2.1). Since larger *β* probes 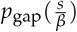 at ever-smaller arguments, only the behavior of *p*_gap_ (Δ) near Δ = 0 matters.

When *p*_gap_ is *regular* (by this I mean 0 < *p*_gap_(0) < ∞), Eq. 15 can be evaluated to leading order:

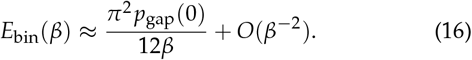

The saddle-point condition then gives 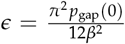, which inverts to 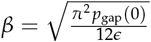. Substituting Eq. 16 and this expression for *β* back into Eq. 13 yields our desired result:

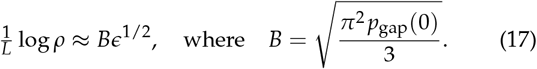

It is tempting to assume that *p*_gap_ is always regular, and indeed it is if each *θ*_*lc*_ parameter is drawn i.i.d. from a sufficiently well-behaved distribution (see SI Sec. S3). Regularity is not guaranteed, however, when the magnitude of additive effects varies across positions. For example, if most positions in a sequence have little to no effect, the gap distribution may scale as Δ^*γ*^ near Δ = 0 for some negative exponent *γ*.^2^ This is an important case to consider because many empirical landscapes exhibit this type of behavior.

We therefore consider the more general case where *p*_z_(Δ)∼ *c* Δ^*γ*^ near Δ = 0, for some exponent *γ* > − 1 and coefficient *c* > 0. Repeating the asymptotic analysis with this assumption yields a more general power law:

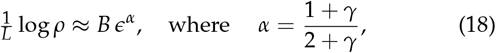

and *B* is a constant that depends on *γ* and *c*; see SI Sec. S2 for a full derivation. Note that the exponent *α* increases monotonically with *γ*. For regular gap distributions (*γ* = 0), one recovers 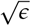 scaling 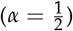. When the gap distribution is enriched for small gaps (*γ* < 0), as is expected when most positions have small fitness effects, the density falls off more quickly near *F*_max_ than 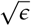 scaling predicts, producing a stubbier peak 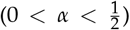. Conversely, a deficit of small gaps (*γ* > 0) causes the density to fall off more slowly, producing a sharper peak 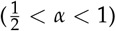.

In the extreme case where all gaps are bounded below by some Δ_0_ > 0, log *ρ* grows approximately linearly with *ϵ* (i.e., *α* = 1), but is also tempered by a log-linear term in *ϵ* that reduces this growth away from *ϵ* = 0. This is exemplified by the landscape proposed by Berg and von Hippel (1987), in which *F*_max_ −*f* (*x*) is Δ_0_ times the Hamming distance between *x* and *x*_max_, and consequently *p*_gap_(Δ) = *δ*(Δ − Δ_0_). See SI Sec. S4 for details.

Figure 3 validates these predictions for binary landscapes with four choices of gap distribution: regular (*γ* = 0), enriched for small gaps (*γ* < 0), depleted of small gaps (*γ* > 0), and fixed gap size (Hamming-distance landscape). In all cases, the near-peak approximation closely tracks *ρ*_saddle_ near *F*_max_, while *ρ*_bulk_ progressively overestimates the density. The same curves plotted against *ϵ*^*α*^ are shown in SI Fig. S1.

**Figure 3.**
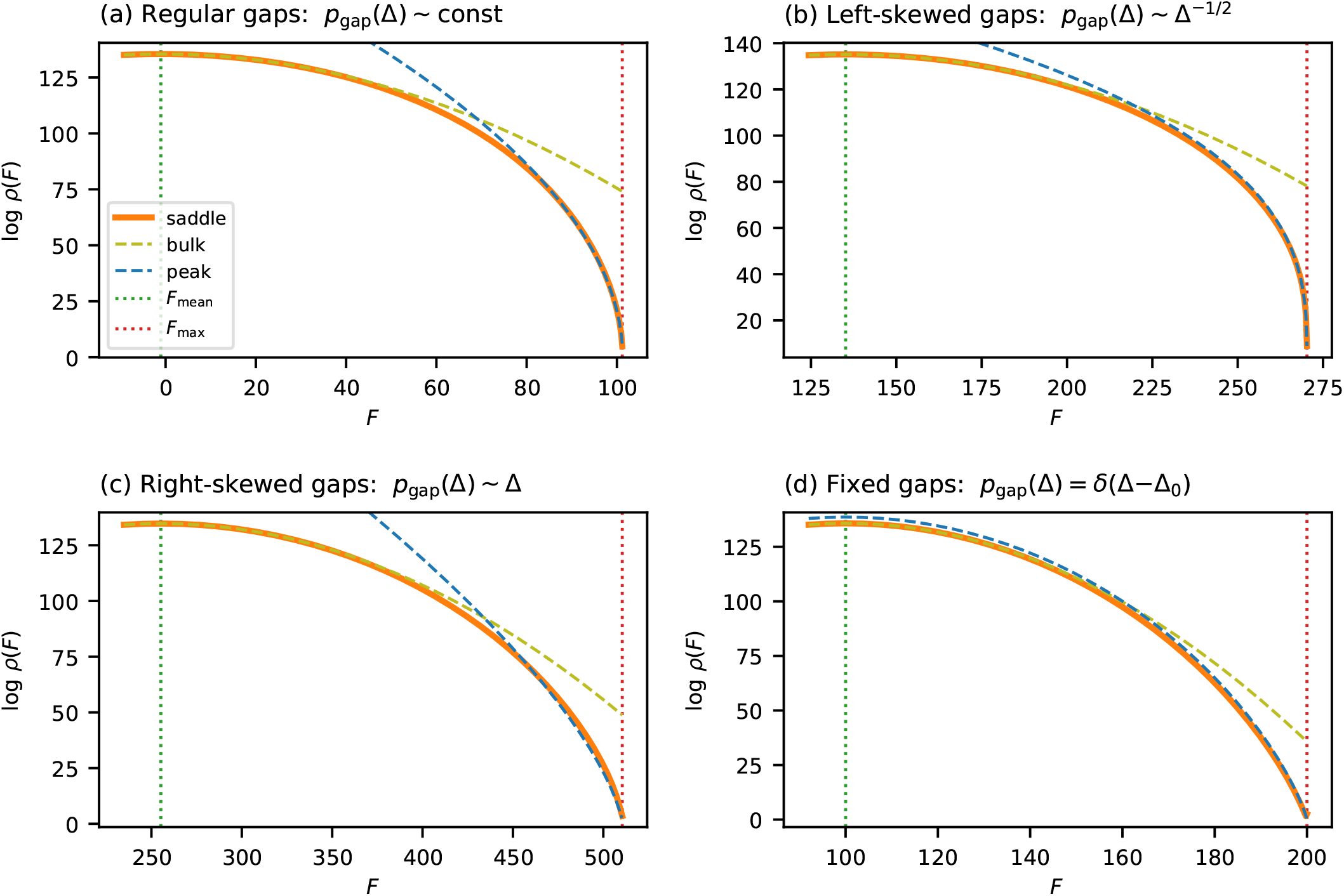
Near-peak scaling for additive landscapes on binary sequences (*C* = 2, *L* = 200). Each panel compares the saddle-point approximation *ρ*_saddle_ (orange solid line) to the Gaussian approximation *ρ*_bulk_ (olive dashed line) and the near-peak approximation *ρ*_peak_ (blue dashed line). (a) Gaussian fitness effects (*θ*_*lc*_ ∼ 𝒩 (0, 1)), producing a regular gap distribution with *γ* = 0 and *α* = 1/2. (b)Left-skewed gap density (*p*_gap_ ∝ Δ^−1/2^, *γ* = − 1/2), enriched for small gaps and producing a stubbier peak (*α* = 1/3). (c) Right-skewed gap density (*p*_gap_ ∝ Δ, *γ* = 1), depleted of small gaps and producing a sharper peak (*α* = 2/3). (d) Hamming-distance landscape with uniform gap Δ = 1 at all positions (*α* = 1 with log-linear correction). Vertical dotted lines indicate *F*_max_ and *F*_mean_. See Fig. S1 for plots made with respect to *ϵ*^*α*^ instead of *F*.

### Near-peak scaling for general alphabets

The preceding analysis was restricted to binary alphabets, where each position contributes a single fitness gap Δ_*l*_ to *E*(*β*). This allowed *E*(*β*) to be evaluated in the large-*L* limit by replacing the sum over positions with an integral over the gap density *p*_gap_(Δ). For alphabets with *C* ≥3 characters, however, *E*(*β*) in Eq. 12 includes contributions from multiple non-optimal characters at each position. The resulting multidimensional integral cannot be evaluated so easily.

Fortunately, we can bound *E*(*β*) both above and below using the binary alphabet results. Define Δ_*l*_ to be the gap between the additive effects of the optimal and runner-up characters at position *l*, and let *E*_bin_(*β*) be the deficit contribution obtained by restricting the alphabet to only these two characters. For the lower bound, this restriction removes terms from the sum inside the logarithm in Eq. 12, which can only decrease *E*(*β*), giving *E*_bin_(*β*) ≤ *E*(*β*). For the upper bound, we use Δ_*l*_ ≤ *δ*_*lc*_ for all 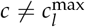, together with the subadditivity of log(1 + *x*), to get

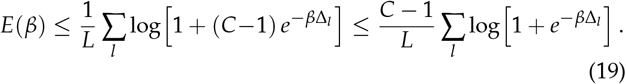

We thus obtain the sandwich bound,

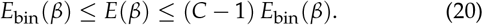

This implies that *E*(*β*) exhibits the same asymptotic behavior in the *β* → ∞ limit as its binary alphabet counterpart. In particular, when the runner-up gap distribution satisfies *p*_gap_(Δ)∼*c* Δ^*γ*^ near the origin, the scaling exponent *α* = (1 + *γ*)/(2 + *γ*) carries over unchanged from the binary case, though the constant *B* may differ.

Accurately describing real-world landscapes requires one additional consideration. If all *C* characters at a position make the same contribution to fitness (*δ*_*lc*_ = 0 for all *c*), that position contributes a constant ^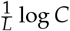^ to *E*(*β*) that is independent of *β*. The cumulative effect of such positions is to add a constant to ^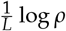^. Similar but smaller contributions arise at positions where some but not all characters are tied for maximal fitness. These considerations motivate the following hypothesis: that the near-peak genotypic density of additive landscapes will often follow an exponentiated power law of the form

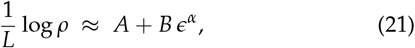

where the specific values of *A, B*, and *α* for each landscape can be determined by fitting this function to the saddle-point approximation in Eq. 11.

I propose Eq. 21 as an effective description of many additive landscapes. I emphasize, though, that this form is not exact and will not accurately describe every landscape. Rather, it is a parametric approximation that is motivated by idealized models and that appears to often work well in practice (including for landscapes with gap distributions that do not follow a power law).

Figure 4 tests this hypothesis on the same four landscapes considered in Fig. 2: two simulated (i.i.d. Gaussian effects with *L* = 100, *C* = 4 and *L* = 100, *C* = 20) and two empirical (*lac* promoter and GB1 protein). For each landscape, I fit *L*^−1^ log *ρ* = *A* + *B ϵ*^*α*^ to the saddle-point density in the near-peak region, with *A, B*, and *α* as free parameters. The two simulated landscapes give *α* ≈ 0.53, close to the theoretical prediction of 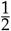 for i.i.d. fitness effects. The two empirical landscapes give *α* ≈ 0.40, slightly below 1/2, consistent with a mild enrichment of small gaps. The same curves plotted against *ϵ*^*α*^, using the fitted *α* value for each panel, are shown in SI Fig. S2.

**Figure 4.**
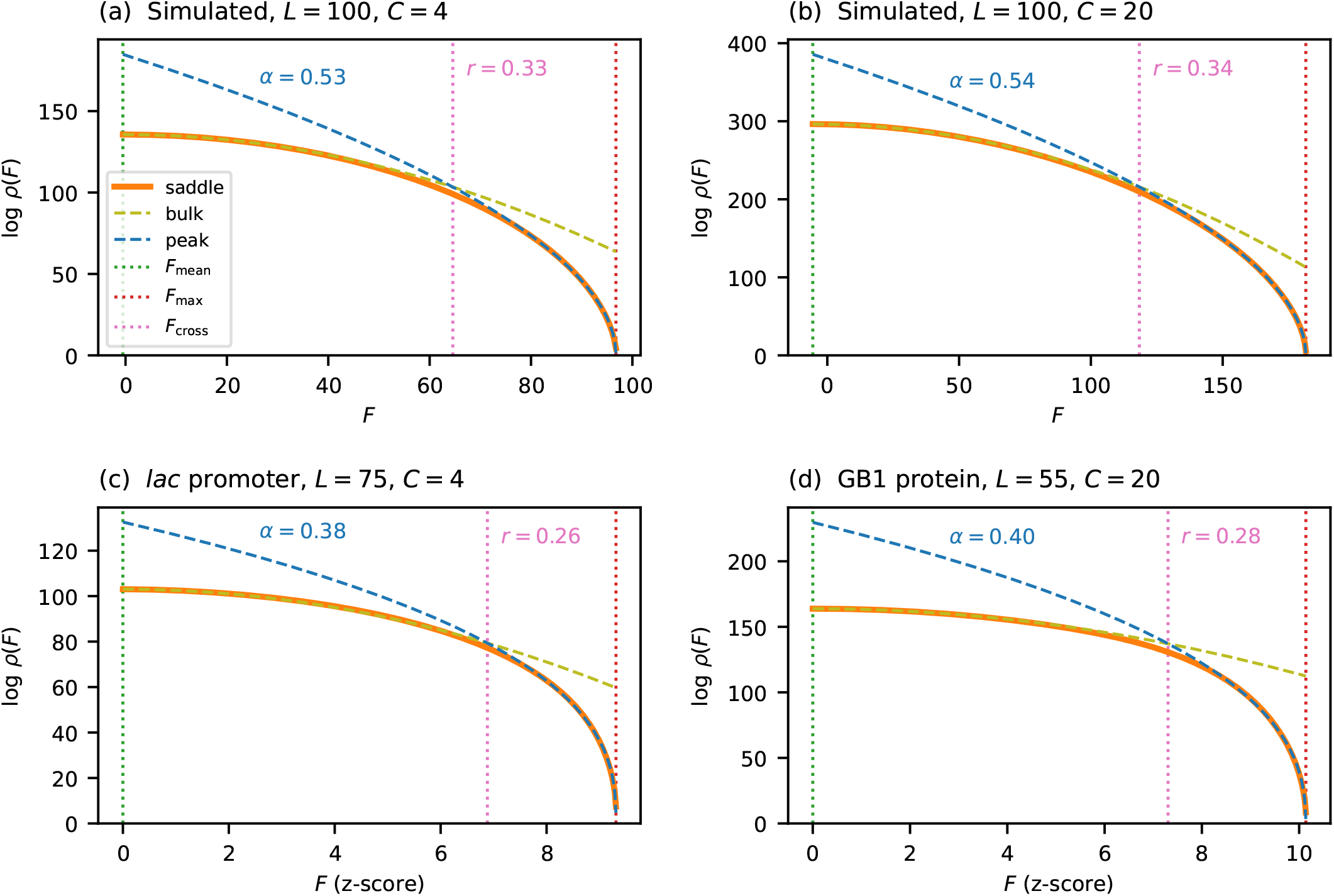
Near-peak scaling for general alphabets, tested on the same four landscapes as in Fig. 2. Each panel plots log *ρ*(*F*) versus *F*, comparing the saddle-point approximation *ρ*_saddle_, the Gaussian approximation *ρ*_bulk_, and the near-peak scaling form *ρ*_peak_, the parameters of which (*A, B*, and *α*) were obtained by fitting *L*^−1^ log *ρ* = *A* + *B ϵ*^*α*^ to each individual landscape. Empirical *F* values are measured in the z-score units defined in Fig. 1. The vertical dotted line at *F*_cross_ marks the crossover fitness at which *ρ*_peak_ becomes a closer approximation to *ρ*_saddle_ than *ρ*_bulk_, with the crossover ratio *r* labeled. (a) Simulated landscape using *C* = 4. (b) Simulated landscape using *C* = 20. (c) *lac* promoter landscape (*L* = 75, *C* = 4). (d) GB1 protein landscape (*L* = 55, *C* = 20). See Fig. S2 for plots made with respect to *ϵ*^*α*^ instead of *F*.

It is useful to ask over what fraction of the fitness range the near-peak scaling outperforms the Gaussian approximation. To quantify this, I define the *crossover fitness F*_cross_ as the fitness value at which *ρ*_peak_ becomes a closer approximation to *ρ*_saddle_ than *ρ*_bulk_ is. The *crossover ratio r* = (*F*_max_−*F*_cross_)/(*F*_max_−*F*_mean_) measures the fraction of the fitness range (from *F*_mean_ to *F*_max_) over which the near-peak scaling dominates. Because both log *ρ*_bulk_ and log *ρ*_peak_ scale linearly with *L*, the fitness at which they cross is determined by an *L*-independent equation, and so the crossover ratio *r* is independent of sequence length. Figure 4 marks *F*_cross_ for each of the four landscapes. Across these landscapes, *r* ranges from 0.26 to 0.34, indicating that the near-peak scaling governs roughly a quarter to a third of the fitness range.

### Single-trait global epistasis models

The results above apply to purely additive landscapes. In many experimental settings, however, the observed fitness is better described not as a direct sum of per-position effects but rather as a nonlinear readout of an underlying additive trait. More precisely, the measured fitness takes the form *F* = *g*(*ϕ*), where *ϕ*(*x*) = *θ*_0_ + ∑_*l,c*_ *θ*_*lc*_ *x*_*lc*_ is an additive trait and *g* is a nonlinear function. Such models are commonly referred to as global epistasis models (Starr and Thornton 2016; Sailer and Harms 2017; Otwinowski *et al*. 2018; Domingo *et al*. 2019).^3^ I now show that, when the fitness maximum occurs at an extremal value of the additive trait and the slope of *g* is nonzero there, the near-peak scaling exponent *α* is preserved, while only the coefficients *A* and *B* are modified.

The key observation is that, sufficiently close to the fitness peak, *g* is well-approximated by its linearization about *ϕ*_max_. Under this linearization, the fitness deficit at each sequence is simply a rescaled version of its trait deficit: 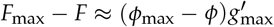, where 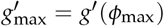. Because this rescaling is linear, it preserves the power-law form of the gap distribution near zero and therefore preserves the scaling exponent *α*.

To state this more precisely, recall that the near-peak scaling of the additive trait (Eq. 21) can be written as

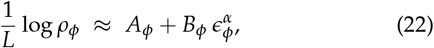

where *ϵ*_*ϕ*_ = (*ϕ*_max_ − *ϕ*)/*L* is the per-site trait deficit. Applying the linearization to transform from trait space to fitness space, and defining the per-site fitness deficit *ϵ*_*F*_ = (*F*_max_ *F*)/*L*, gives (see SI Sec. S5 for derivation):

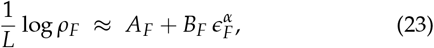

where 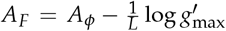 and 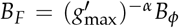. This approximation holds as long as the linearization of *g* remains accurate over the fitness range of interest. In practice, the flatter the nonlinearity is at *ϕ*_max_, the more heavily it compresses the range over which near-peak scaling applies.

Figure 5 illustrates these predictions on a simulated landscape (*L* = 100, *C* = 4) passed through sigmoid nonlinearities of varying steepness. The coefficients *A*_*F*_ and *B*_*F*_ are computed analytically from the scaling behavior of the additive trait *ϕ* and from the value of 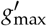, with no additional fitting. In all cases the predicted 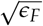scaling closely matches the exact transformed density, confirming that the near-peak exponent is preserved under global epistasis. As expected, sigmoids that are flatter at *ϕ*_max_ (due to greater saturation) yield a larger *B*_*F*_ and a more compressed range of validity.

**Figure 5.**
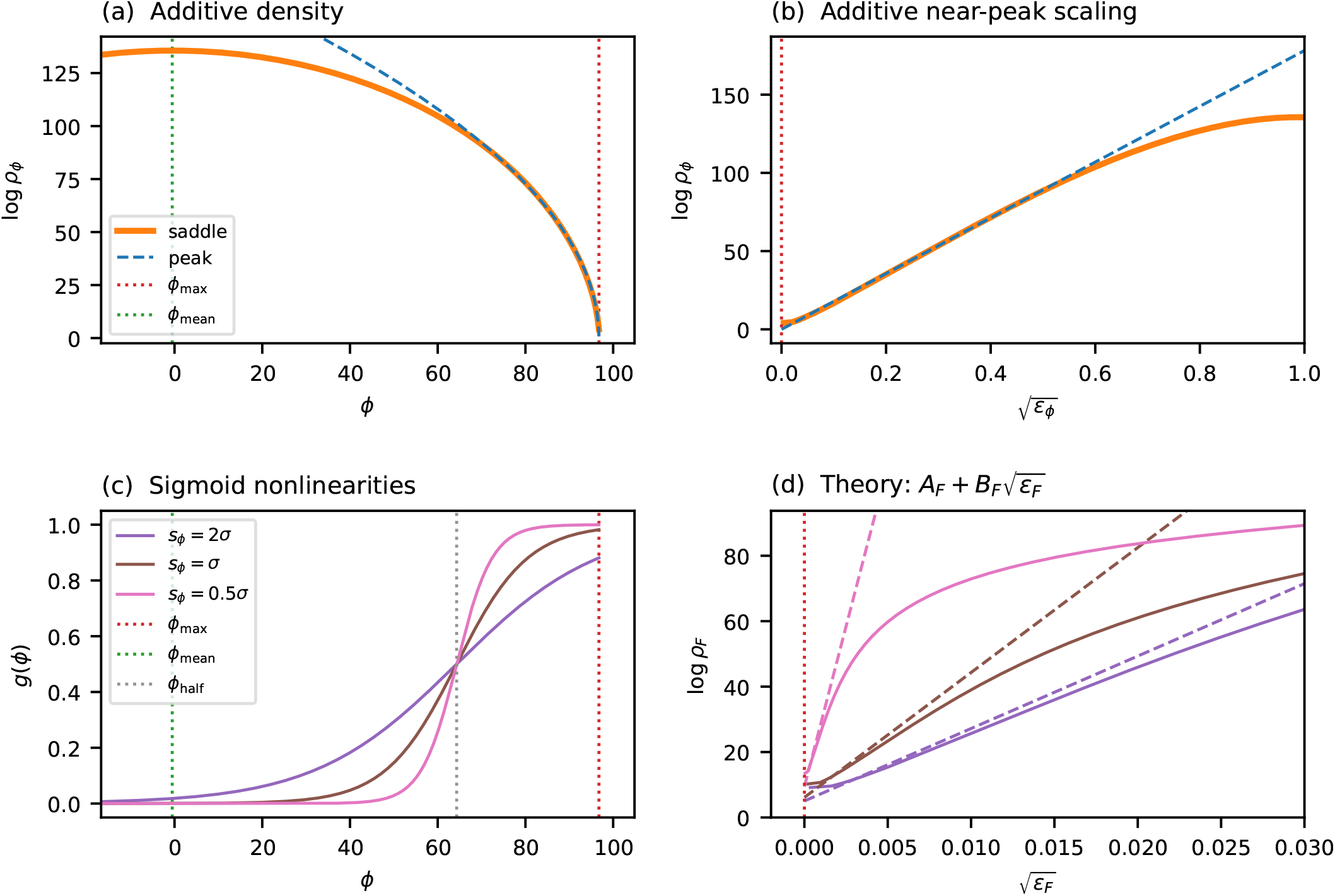
Scaling under global epistasis. A simulated additive landscape (*L* = 100, *C* = 4, Gaussian *θ*_*lc*_) for a trait *ϕ* is transformed by sigmoid nonlinearities *g*(*ϕ*) of varying steepness. (a) The log genotypic density log *ρ*_*ϕ*_ in trait space versus *ϕ*, showing the saddle-point approximation (orange solid line) and the near-peak approximation (blue dashed line). Vertical dotted lines indicate *ϕ*_max_ (red) and *ϕ*_mean_ (green). (b) The same quantities plotted against 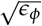, showing the region of approximate linearity from which *B* is extracted. (c) Three sigmoid nonlinearities with midpoint set to 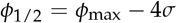 (gray dotted line) and varying span parameter *s*_*ϕ*_, which controls steepness. (d) The transformed density log *ρ*_*F*_ versus 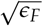for each sigmoid (solid lines) compared to the theoretical prediction 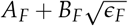(dashed lines in matching colors), where *A*_*F*_ and *B*_*F*_ are computed analytically from *A*_*ϕ*_, *B*_*ϕ*_, and *g*^′^_max_ with no additional fitting (see SI Sec. S5).

### Multi-trait global epistasis models

The preceding section considered fitness landscapes defined by a nonlinear function of a single additive trait. More generally, however, fitness may depend on multiple additive traits simultaneously. I now show that the saddle-point machinery and the near-peak scaling results derived above extend naturally to this multi-trait setting. Consider *K* additive traits

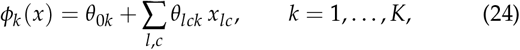

Taken together, these map each sequence *x* to a *K*-dimensional trait vector 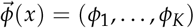. To define a tilted distribution, we introduce a complementary vector of *K* inverse temperatures 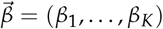. The resulting tilted distribution has the form 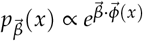. The key observation is that the exponent 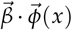 is itself an additive function of *x*, with effective per-position parameters 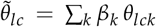. The tilted distribution therefore factorizes across positions exactly as in the single-trait case. The corresponding cumulant-generating function is

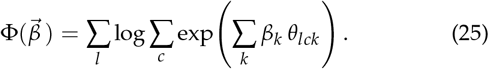

Its gradient ∇Φ gives the mean of each trait under the tilted distribution, and the entries of the Hessian matrix Φ^′′^ give the covariances.

The saddle-point approximation generalizes directly (Barndorff-Nielsen and Cox 1989, Sec. 6.5). As in the single-trait case, we choose 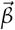 so that the target trait vector 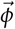coincides with the mean of the tilted distribution. Because Φ is strictly convex, this condition uniquely determines 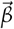. The resulting genotypic density, which is joint across *K* traits, is

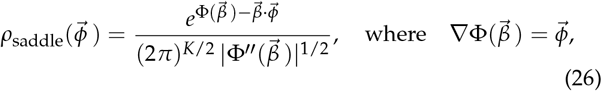

and where | Φ^′′^| denotes the determinant of the Hessian. This reduces to Eq. 11 when *K* = 1.

The near-peak scaling behavior of single-trait global epistasis models also extends to the multi-trait setting. Suppose fitness is a nonlinear function of the trait vector,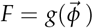. The achievable trait vectors form a bounded region in *K*-dimensional trait space. If *g* attains its maximum *F*_max_ on the boundary of this region and has nonzero gradient there, then *g* can be linearized in the vicinity of the maximum. This linearization projects the *K* traits onto a single effective additive trait,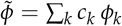, where 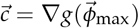. The near-peak scaling results from the preceding sections then apply directly, with the exponent *α* determined by the gap distribution of 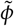. As in the single-trait case, the range of fitness values over which this scaling holds is curtailed by the nonlinearity of *g*.

## Discussion

The core finding of this work is that the log genotypic density of additive fitness landscapes typically follows a power law near maximal fitness (Eq. 21). The exponent in the power law, *α*, determines the shape of this density near the peak: *α* is constrained to lie between 0 and 1, with larger *α* producing sharper peaks and smaller *α* producing stubbier ones. The value of *α* depends on how the fitness gaps between optimal and runner-up characters at each position are distributed, and thus varies from landscape to landscape. Notably, *α* is independent of both sequence length and alphabet size. In idealized landscapes, where the additive effects are drawn from a single distribution, one obtains 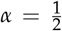 regardless of the specific distribution used. In more realistic situations, however, the scale of additive effects can vary substantially from position to position, causing *α* to deviate from 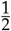. In particular, one should often expect to find 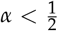 (increased stubbiness) on empirical landscapes because positions with small-to-negligible effects are often much more frequent than positions with large effects.

These results, to my knowledge, provide the first quantitative description of near-peak abundance of genotypes for any class of fitness landscapes.^4^ Indeed, the Gaussian approximation suggested by the CLT fundamentally breaks down near the fitness maximum of additive landscapes. If taken literally, it predicts that many sequences lie far above *F*_max_. If the Gaussian approximation is instead truncated at maximal fitness, it still predicts a vast overabundance of high-fitness sequences, often yielding densities that are off by many orders of magnitude. By contrast, the exponentiated power law that I describe is highly accurate in this regime. It reveals that, as fitness is decreased from its maximum, the number of available genotypes initially expands at a near-infinite rate, then grows increasingly slowly until it settles down to the growth rate predicted by the Gaussian approximation. The result is a broad and gently rounded (i.e., stubby) summit.

The exponentiated power law given in Eq. 21 comes with several caveats. First, it applies only to additive and globally epistatic landscapes. In particular, the consequences of specific epistatic interactions on near-peak density have yet to be investigated. Second, the derivation assumes that sequence length *L* is large. In practice, however, this scaling behavior is often observed even for landscapes describing sequences of length *L*∼ 10, such as TF binding motifs; see Fig. S3. Third, the derivation assumes that the distribution of position-specific fitness gaps behaves as a power law near zero. That said, Eq. 21 appears to provide a good empirical fit to many additive landscapes, even when *α, A*, and *B* must be fit empirically rather than derived from the gap distribution.

The scaling behavior also extends to globally epistatic landscapes defined on one or more additive traits. This only happens, however, under certain conditions: the nonlinearity must attain its maximum value on the boundary of the accessible trait space, and it must do so with nonzero gradient. If these conditions are satisfied, the nonlinearity can be approximated by a linear function near the peak, and the problem of computing genotypic densities reduces to that of a one-dimensional additive fitness function. If there is just one additive trait, the near-peak density will exhibit the same exponent *α* (and thus the same “stubbiness”) as the additive trait itself. When there are multiple additive traits, the linearized fitness landscape will be a linear combination of the traits, and is therefore expected to exhibit an empirical *α* intermediate to the *α* values for the individual traits (the math here is not so clean, though). An important caveat with all such global epistasis models is that the scaling behavior will only be valid over the range of *F* values for which linearization provides a good approximation. This range will be particularly small for global epistasis nonlinearities that plateau near their maxima.

Looking forward, the mathematical methods used in this work might prove useful for addressing other questions about fitness landscapes. The distribution of fitness effects (DFE) is of central importance in evolutionary genetics (Gillespie 1983; Eyre-Walker and Keightley 2007), and future work using related mathematical approaches could potentially be used to study the DFE near fitness peaks. Related questions, such as the distribution of Hamming distances among sequences at a given fitness level, might also be addressed. Finally, the multi-trait saddle-point approximation in Eq. 26 naturally suggests applications to Fisher’s geometric model (Fisher 1930; Orr 1998). It would be particularly interesting, in the spirit of Hwang *et al*. (2017), to see if similar techniques could shed light on how the dramatic variation in genotypic density that occurs across the fitness range affects the predictions of Fisher’s model.

## Acknowledgments

This project grew out of conversations with Ville Mustonen and Michael Lässig back in 2006. I also thank my colleagues David McCandlish and Peter Koo for helpful discussions. Financial support was provided by NIH grants R01HG011787 and S10OD028632.

## Data availability

All the code used to carry out this analysis and generate the figures is publicly available at https://github.com/jbkinney/26_stubby.

## Supplemental Information

Additional mathematical details are provided in Supplemental Information.

## Footnotes

1 I use the term *fitness* to refer to any scalar function of sequence, regardless of whether that function represents evolutionary pressure, biological activity, or some biophysical quantity.

2 We require *γ* > −1 in order for *p*_gap_ to be integrable near Δ = 0.

3 Note that this definition of global epistasis is based on the structure of the underlying equations and is distinct from the phenomenological definition sometimes used in discussions of diminishing returns epistasis, e.g., (Kryazhimskiy *et al*. 2014; Reddy and Desai 2021).

4 That is, apart from the highly idealized Hamming-distance landscape studied by Berg and von Hippel (1987).

