## Supplemental Information for "Mount Fuji’s stubby peak: the genotypic density of additive landscapes near maximal fitness"

Justin B. Kinney

May 6, 2026

**S4 Near-peak scaling for the Hamming-distance landscape** **6**

**S5 Global epistasis coefficients** **6**

#### S1 Cumulants of the fitness distribution for additive landscapes

Using the expression for $\Phi(\beta)$ in Eq. 7 of the main text and the fact that $\Phi$ is a cumulant generating function, the mean and variance of $p_\beta(F)$ are found to be:

$$\mu_\beta = \Phi'(\beta) = \theta_0 + \sum_l \sum_c w_{lc}\,\theta_{lc}, \tag{S1}$$

$$\sigma_\beta^2 = \Phi''(\beta) = \sum_l \left[ \sum_c w_{lc}\,\theta_{lc}^2 - \left( \sum_c w_{lc}\,\theta_{lc} \right)^2 \right], \tag{S2}$$

where the weights $w_{lc}$ are defined to be

$$w_{lc} = \frac{e^{\beta\theta_{lc}}}{\sum_{c'} e^{\beta\theta_{lc'}}}. \tag{S3}$$

More generally, the $n$th cumulant is given by $\kappa_n = \Phi^{(n)}(\beta)$. $\Phi$ scales linearly with $L$, and so do all these cumulants, including $\sigma_\beta^2$. The standardized cumulants therefore scale as $\kappa_n/\sigma_\beta^n = O(L^{1-n/2})$,

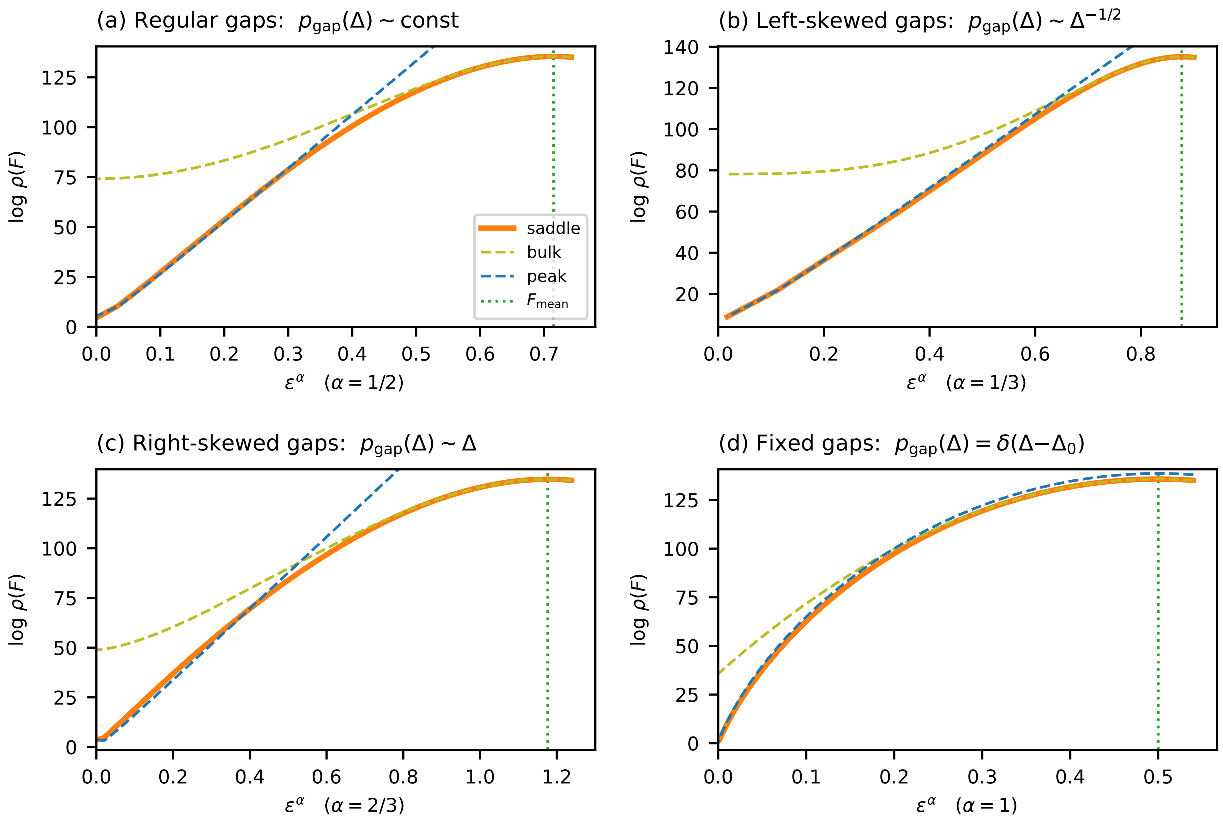

Figure S1: Near-peak scaling for binary additive landscapes plotted against $\epsilon^\alpha$, where $\epsilon = (F_{\max} - F)/L$ and $\alpha$ is the analytic near-peak exponent for each panel. Curves are the same as in Fig. 3.

and thus vanish as $L \to \infty$ for all $n \geq 3$. This confirms that the tilted fitness distribution converges to a Gaussian in the large $L$ limit (Eq. 8 of the main text).

### S2 Supporting derivations for the near-peak scaling

This section provides mathematical details supporting the near-peak scaling results in Eqs. 17−18 of the main text.

#### S2.1 Evaluation of $\int_0^\infty \log(1 + e^{-u})\, du$

The integral $\int_0^\infty \log(1 + e^{-u})\, du$ appearing in Eq. 16 of the main text is evaluated by expanding the logarithm as a series:

$$\int_0^\infty \log(1 + e^{-u})\, du = \sum_{k=1}^{\infty} \frac{(-1)^{k+1}}{k} \int_0^\infty e^{-ku}\, du = \sum_{k=1}^{\infty} \frac{(-1)^{k+1}}{k^2} = \eta(2), \tag{S4}$$

where $\eta(s) = \sum_{k=1}^{\infty} (-1)^{k+1}/k^s$ is the Dirichlet eta function. To evaluate $\eta(s)$, we split the sum into odd and even terms:

$$\eta(s) = \sum_{k=1}^{\infty} \frac{1}{k^s} - 2 \sum_{k=1}^{\infty} \frac{1}{(2k)^s} = \zeta(s) - 2^{1-s}\, \zeta(s) = (1 - 2^{1-s})\, \zeta(s), \tag{S5}$$

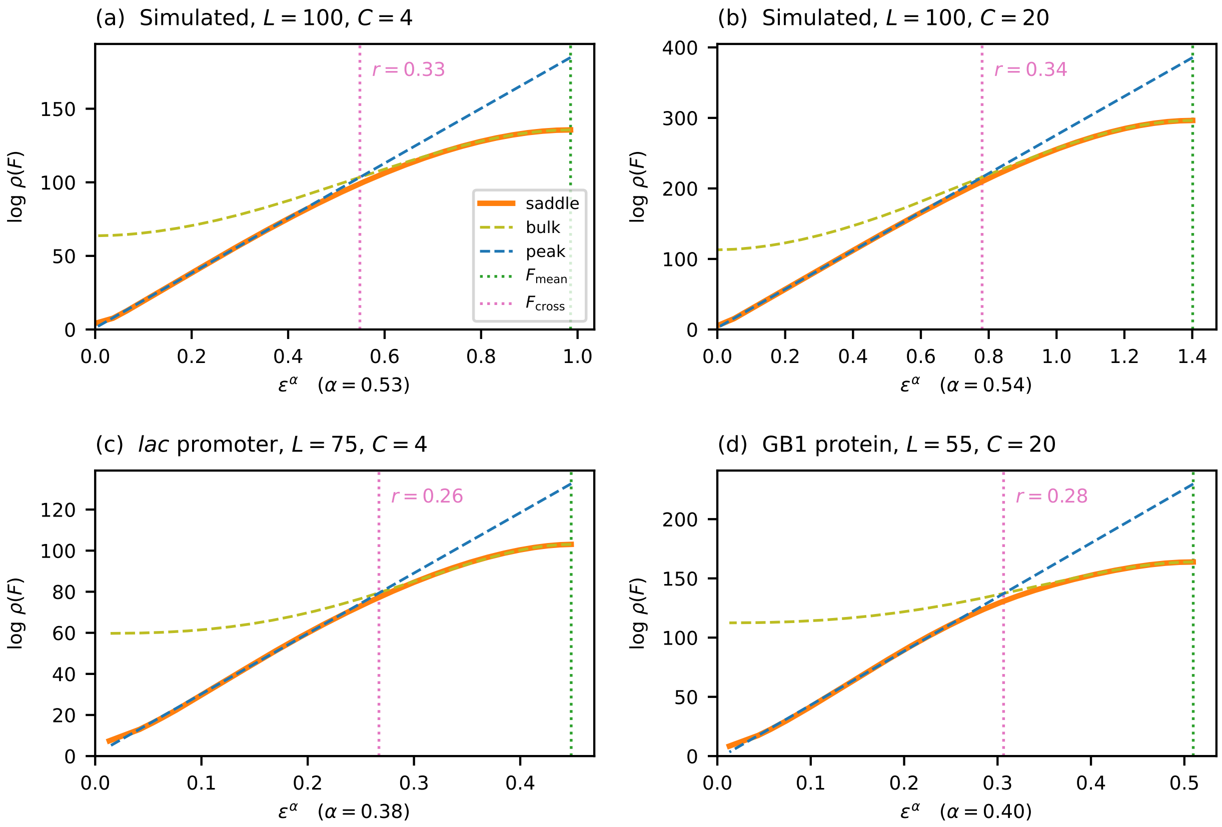

Figure S2: Near-peak scaling for general alphabets plotted against $\epsilon^\alpha$, where $\alpha$ is fit separately for each panel. Curves are the same as in Fig. 4.

where $\zeta(s) = \sum_{k=1}^{\infty} 1/k^s$ is the Riemann zeta function. Using the Basel identity $\zeta(2) = \pi^2/6$, we obtain $\eta(2) = (1 - 2^{-1}) \cdot \pi^2/6 = \pi^2/12$, and thus

$$\int_0^\infty \log(1 + e^{-u})\, du = \frac{\pi^2}{12}. \tag{S6}$$

#### S2.2 Generalized integral identity

The derivation in Sec. S2.3 below relies on the identity

$$I(\gamma) \equiv \int_0^\infty \log(1 + e^{-u})\, u^\gamma\, du = \Gamma(1+\gamma)\, \eta(2+\gamma), \qquad \gamma > -1. \tag{S7}$$

To derive this identity, expand $\log(1 + e^{-u}) = \sum_{k=1}^{\infty} (-1)^{k+1} e^{-ku}/k$ and integrate term by term:

$$I(\gamma) = \sum_{k=1}^{\infty} \frac{(-1)^{k+1}}{k} \int_0^\infty u^\gamma\, e^{-ku}\, du = \sum_{k=1}^{\infty} \frac{(-1)^{k+1}\, \Gamma(1+\gamma)}{k^{2+\gamma}} = \Gamma(1+\gamma)\, \eta(2+\gamma). \tag{S8}$$

As a consistency check, setting $\gamma = 0$ gives $I(0) = \Gamma(1)\, \eta(2) = \pi^2/12$, recovering Eq. S4.

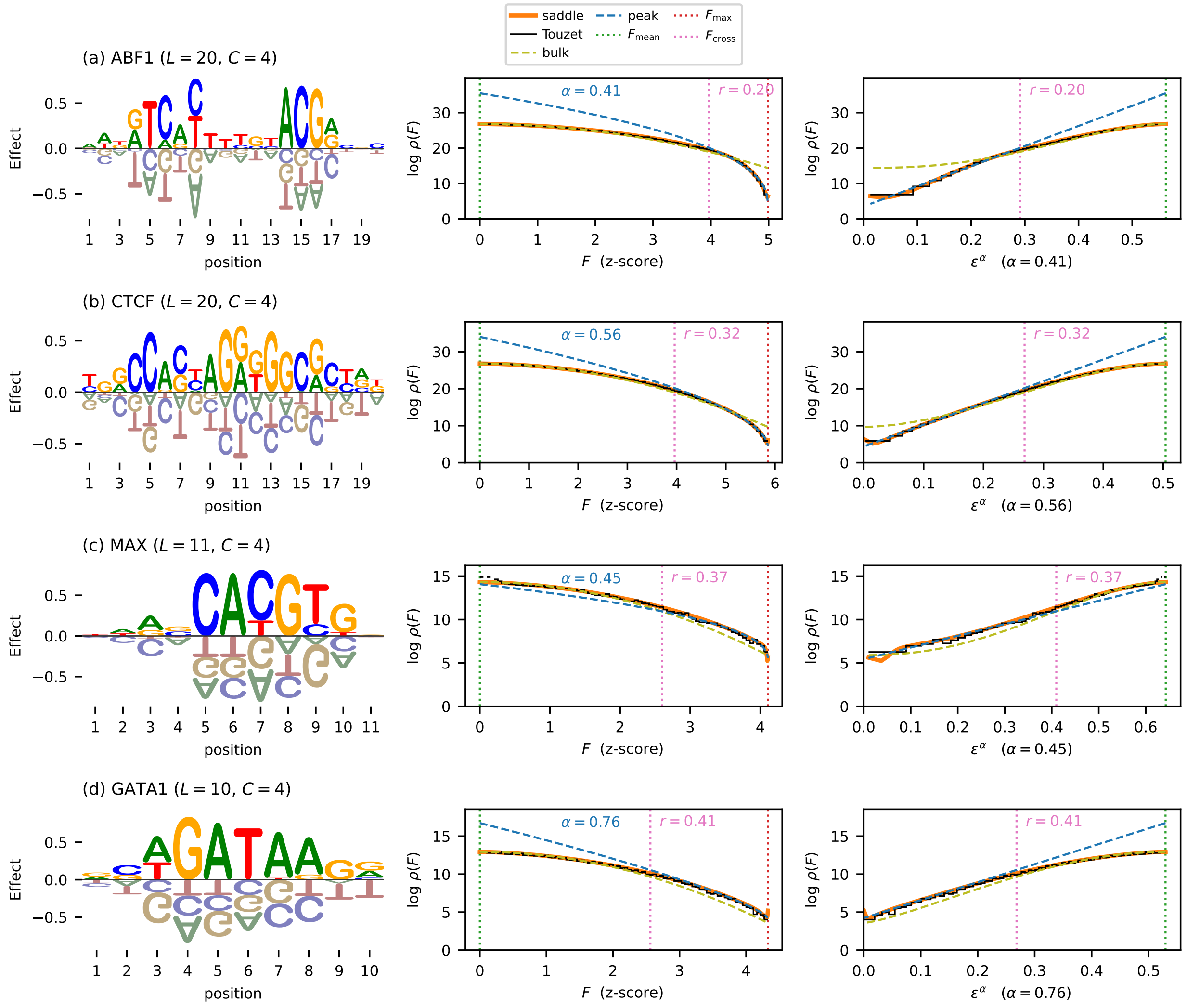

Figure S3: Near-peak genotypic densities for four transcription factors (TFs). (a) The ABF1 motif reported by Mustonen et al. [1]. (b-d) The CTCF, MAX, and GATA1 motifs from HOCOMOCO v14 [2]. Left panels show additive effects shifted and scaled so that $F$ values correspond to z-scores [$F_{\mathrm{mean}} = 0$, $\mathrm{var}(F) = 1$]. Middle panels compare the saddle-point approximation, Touzet density estimate [3], Gaussian bulk approximation, and fitted near-peak scaling behavior (i.e., $A + B\epsilon^\alpha$) as functions of $F$. Right panels show the same curves as functions of $\epsilon^\alpha$. The values of $A$, $B$, and $\alpha$ used in each row were empirically determined from the corresponding saddle-point approximation.

#### S2.3 Derivation of the generalized scaling exponent

When the gap distribution behaves as $p_{\mathrm{gap}}(\Delta) \sim c\,\Delta^\gamma$ near $\Delta = 0$ with $\gamma > -1$, the substitution $u = \beta\Delta$ in the integral form of $E_{\mathrm{bin}}(\beta)$ (Eq. 15 of the main text) gives

$$E_{\mathrm{bin}}(\beta) \;\sim\; \frac{c}{\beta^{1+\gamma}} \int_0^\infty \log(1 + e^{-u})\, u^\gamma \, du \;=\; \frac{b}{\beta^{1+\gamma}}, \tag{S9}$$

where $b = c\,\Gamma(1+\gamma)\,\eta(2+\gamma)$ using the identity in Eq. S7. The saddle-point equation $\epsilon = -E'_{\text{bin}}(\beta)$ then gives

$$\epsilon = \frac{(1+\gamma)\,b}{\beta^{2+\gamma}}, \qquad \text{so} \qquad \beta = \left[\frac{(1+\gamma)\,b}{\epsilon}\right]^{1/(2+\gamma)}. \tag{S10}$$

From Eq. 13 of the main text, $\frac{1}{L}\log\rho_{\text{saddle}} \approx E_{\text{bin}}(\beta) + \beta\epsilon$. Combining these two terms:

$$E_{\text{bin}}(\beta) + \beta\epsilon = \frac{b}{\beta^{1+\gamma}} + \beta\epsilon = \frac{\epsilon\,\beta}{1+\gamma} + \beta\epsilon = \frac{(2+\gamma)}{(1+\gamma)}\,\beta\epsilon. \tag{S11}$$

Substituting the expression for $\beta$ yields

$$\frac{1}{L}\log\rho \;\sim\; \underbrace{\frac{(2+\gamma)}{(1+\gamma)}\big[(1+\gamma)\,b\big]^{1/(2+\gamma)}}_{B}\;\epsilon^{\overbrace{(1+\gamma)/(2+\gamma)}^{\alpha}}. \tag{S12}$$

This yields the scaling exponent $\alpha = (1+\gamma)/(2+\gamma)$ and the coefficient $B$ stated in Eq. 18 of the main text.

### S3 Gap density for continuous distributions

The main text states that the condition $0 < p_{\text{gap}}(0) < \infty$ is always satisfied when the fitness effects $\theta_{lc}$ are independently drawn from the same continuous distribution. Here we prove this claim and evaluate $p_{\text{gap}}(0)$ for the Gaussian distribution used in simulations.

#### S3.1 General formula from order statistics

Let $\theta_{(1)} \geq \theta_{(2)} \geq \cdots \geq \theta_{(C)}$ denote the order statistics of $C$ i.i.d. draws from a density $p_\theta$ with CDF $P_\theta$. The gap $\Delta = \theta_{(1)} - \theta_{(2)}$ between the largest and second-largest values has density

$$p_{\text{gap}}(\delta) = C(C-1)\int_{-\infty}^{\infty} p_\theta(m+\delta)\,p_\theta(m)\,P_\theta(m)^{C-2}\,dm, \qquad \delta \geq 0, \tag{S13}$$

which follows from the joint density of the top two order statistics, $f_{\theta_{(1)},\theta_{(2)}}(x_1,x_2) = C(C-1)\,p_\theta(x_1)\,p_\theta(x_2)\,P_\theta(x_2)^{C-2}$ for $x_1 \geq x_2$, by substituting $\delta = x_1 - x_2$ and integrating over $m = x_2$. Setting $\delta = 0$ gives

$$p_{\text{gap}}(0) = C(C-1)\int_{-\infty}^{\infty} p_\theta(m)^2\,P_\theta(m)^{C-2}\,dm. \tag{S14}$$

We now show that $0 < p_{\text{gap}}(0) < \infty$ for any continuous density $p_\theta$. The integrand $C(C-1)\,p_\theta(m)^2\,P_\theta(m)^{C-2}$ is non-negative everywhere and strictly positive on the interior of the support of $p_\theta$ (where $p_\theta(m) > 0$ and $0 < P_\theta(m) < 1$). It follows that $p_{\text{gap}}(0) > 0$. For the upper bound, $p_\theta(m)^2 \leq \|p_\theta\|_\infty\,p_\theta(m)$ and $P_\theta(m)^{C-2} \leq 1$, so $p_{\text{gap}}(0) \leq C(C-1)\,\|p_\theta\|_\infty < \infty$.

#### S3.2 Evaluation for Gaussian effects

For $\theta_{lc} \sim \mathcal{N}(0, s^2)$, substituting $p_\theta(m) = \varphi(m/s)/s$ and $P_\theta(m) = \Phi_{\text{N}}(m/s)$ into Eq. S14 and changing variables to $z = m/s$ gives

$$p_{\text{gap}}(0) = \frac{C(C-1)}{s}\int_{-\infty}^{\infty} \varphi(z)^2\,\Phi_{\text{N}}(z)^{C-2}\,dz, \tag{S15}$$

where $\varphi$ and $\Phi_N$ denote the standard normal density and CDF, respectively. For $C = 2$ the integral evaluates to $1/(2\sqrt{\pi})$, giving $p_{\text{gap}}(0) = 1/(\sqrt{\pi}\, s)$. For general $C$ the integral must be computed numerically; the resulting dimensionless product $s \cdot p_{\text{gap}}(0)$ is tabulated below for the alphabet sizes used in the main text.

| $C$ | $s \cdot p_{\text{gap}}(0)$ |
|---|---|
| 2 | 0.564 |
| 4 | 1.029 |
| 20 | 1.867 |

#### S4 Near-peak scaling for the Hamming-distance landscape

The Hamming-distance landscape [4] assumes that every mutation away from the optimal sequence $x_{\max}$ incurs the same fitness cost $\Delta_0 > 0$. A sequence at Hamming distance $k$ from $x_{\max}$ therefore has fitness $F_{\max} - k\Delta_0$. Because there are $\binom{L}{k}(C-1)^k$ such sequences, the exact genotypic density is

$$\rho(F_{\max} - k\Delta_0) = \binom{L}{k}(C-1)^k, \qquad k = 0, 1, \ldots, L. \tag{S16}$$

For small per-site fitness deficit $\epsilon = k\Delta_0/L$ with $k \ll L$, Stirling's approximation $\log\binom{L}{k} \approx k\log(eL/k)$ gives

$$\frac{1}{L}\log\rho(F_{\max} - L\epsilon) \approx \frac{\epsilon}{\Delta_0}\log\frac{e(C-1)\Delta_0}{\epsilon}. \tag{S17}$$

For $C = 2$ this reduces to $(\epsilon/\Delta_0)\log(e\Delta_0/\epsilon)$.

This scaling is qualitatively different from the power-law form $\log\rho \sim \epsilon^\alpha$ derived in the main text. The difference is traceable to the gap distribution: in the Hamming-distance landscape every position has the same gap $\Delta_0 > 0$, so $p_{\text{gap}}(\Delta) = \delta(\Delta - \Delta_0)$. This is not consistent with the assumption $p_{\text{gap}}(\Delta) \sim c\,\Delta^\gamma$ near $\Delta = 0$ for any $c$ and $\gamma$, so the power-law scaling does not apply.

The same conclusion follows from the tilting framework. The cumulant generating function is $\Phi(\beta) = L\log[1 + (C-1)e^{-\beta\Delta_0}]$, and the saddle-point condition gives

$$\epsilon = (C-1)\Delta_0\,\frac{e^{-\beta\Delta_0}}{1 + (C-1)e^{-\beta\Delta_0}} \;\approx\; (C-1)\Delta_0\, e^{-\beta\Delta_0} \tag{S18}$$

for large $\beta$. The fitness deficit decreases exponentially in $\beta$, rather than as a power law of $\beta$ as in the generic case (Eq. S9), confirming the breakdown of the power-law scaling mechanism.

#### S5 Global epistasis coefficients

Here we derive the coefficients $A_F$ and $B_F$ appearing in Eq. 23 of the main text. The observed fitness is $F = g(\phi)$, where $g$ is a nonlinear function of the additive trait $\phi$. We assume that $g$ attains its maximum at $\phi_{\max}$ and that $g'_{\max} \equiv g'(\phi_{\max}) \neq 0$, so that $g$ is locally invertible near the peak. In this region the genotypic density transforms by the standard change of variables:

$$\rho_F(F) = \frac{\rho_\phi(\phi)}{g'(\phi)}, \qquad \phi = g^{-1}(F). \tag{S19}$$

Define per-site trait and fitness deficits $\epsilon_\phi = (\phi_{\max} - \phi)/L$ and $\epsilon_F = (F_{\max} - F)/L$. Taylor-expanding $g(\phi_{\max} - L\epsilon_\phi)$ around $\phi_{\max}$ gives

$$\epsilon_F = g'_{\max}\,\epsilon_\phi + O(\epsilon_\phi^2), \tag{S20}$$

so that $\epsilon_\phi \approx \epsilon_F / g'_{\max}$ when $\epsilon_\phi \ll 2g'_{\max}/|g''_{\max}|$. Taking logarithms of Eq. S19 and dividing by $L$:

$$\frac{1}{L}\log\rho_F = \frac{1}{L}\log\rho_\phi - \frac{1}{L}\log g'(\phi). \tag{S21}$$

Near the peak, $g'(\phi) \approx g'_{\max}$, so the second term contributes $-\frac{1}{L}\log g'_{\max}$. Using Eq. 22 of the main text, $L^{-1}\log\rho_\phi \approx A_\phi + B_\phi\,\epsilon_\phi^\alpha$, and substituting $\epsilon_\phi \approx \epsilon_F/g'_{\max}$:

$$\frac{1}{L}\log\rho_F \;\approx\; \underbrace{\left(A_\phi - \frac{\log g'_{\max}}{L}\right)}_{A_F} + \underbrace{\frac{B_\phi}{(g'_{\max})^\alpha}}_{B_F}\,\epsilon_F^\alpha. \tag{S22}$$

The scaling exponent $\alpha$ is unchanged. This approximation is valid when $\epsilon_F \ll 2(g'_{\max})^2/|g''_{\max}|$, i.e., when the nonlinearity is well-approximated by its local linearization near the peak.
